# Dynamic information routing in complex networks

**DOI:** 10.1101/029405

**Authors:** Christoph Kirst, Marc Timme, Demian Battaglia

## Abstract

Flexible information routing fundamentally underlies the function of many biological and artificial networks. Yet, how such systems may specifically communicate and dynamically route information is not well understood. Here we identify a generic mechanism to route information on top of collective dynamical reference states in complex networks. Switching between collective dynamics induces flexible reorganization of information sharing and routing patterns, as quantified by delayed mutual information and transfer entropy measures between activities of a network's units. We demonstrate the power of this generic mechanism specifically for oscillatory dynamics and analyze how individual unit properties, the network topology and external inputs coact to systematically organize information routing. For multi-scale, modular architectures, we resolve routing patterns at all levels. Interestingly, local interventions within one sub-network may remotely determine non-local network-wide communication. These results help understanding and designing information routing patterns across systems where collective dynamics co-occurs with a communication function.

---

Attuned function of many biological or technological networks relies on the precise yet dynamic communication between their subsystems. For instance, the behavior of cells depends on the coordinated information transfer within gene regulatory networks [1, 2] and flexible integration of information is conveyed by the activity of several neural populations during brain function [3]. Identifying general mechanisms for the routing of information across complex networks thus constitutes a key theoretical challenge with applications across fields, from systems biology to the engineering of smart distributed technology [4–6].

Complex systems with a communication function often show characteristic dynamics. such as oscillatory or synchronous collective dynamics with a stochastic component [7–11]. Information is carried in the presence of these dynamics within and between neural circuits [12, 13], living cells [14, 15], ecologic or social groups [16, 17] as well as technical communication systems, such as ad hoc sensor networks [18, 19]. While such dynamics could simply reflect the properties of the interacting unit’s, emergent collective dynamical states in biological networks can actually contribute to the system’s function. For example, it has been hypothesized that the widely observed oscillatory phenomena in biological networks enable emergent and flexible information routing [12].

Here we derive a theory that shows how information if conveyed by fluctuations around collective dynamical reference states (e.g. a stable oscillatory pattern) can be flexibly routed across complex network topologies. Quantifying information sharing and transfer by time-delayed mutual information [20, 21] and transfer entropy [22] curves between time-series of the network’s units, we demonstrate how switching between multi-stable states enables the rerouting of information without any physical changes to the network. In fully symmetric networks, anisotropic information transfer can arise via symmetry breaking of the reference dynamics. For networks of coupled oscillators our approach gives analytic predictions how the physical coupling structure, the oscillators’ properties and the dynamical state of the network co-act to produce a specific communication pattern. Resorting to a collective-phase description [23], our theory further resolves communication patterns at all levels of multi-scale, modular topologies [24, 25], as ubiquitous, e.g., in the brain connectome and bio-chemical regulatory networks [26–29]. We thereby uncover how local interventions within one module may remotely modify information sharing and transfer between other distant sub-networks. A combinatorial number of information routing patterns in networks emerge due to switching between multi-stable dynamical states that are localized on individual subsets of network nodes.

These results offer a generic mechanism for self-organized and flexible information routing in complex networked systems. For oscillatory dynamics the links made between multi-scale connectivity, collective network dynamics and flexible information routing has potential applications to the reconstruction and design of gene regulatory circuits [15, 30], wireless communication networks [4, 19] or to the analysis of cognitive functions [31–35], among others.

## I. INFORMATION ROUTING VIA COLLECTIVE DYNAMICS

To understand how bits of information from external or locally computed signals can be specifically distributed through a network or to it’s downstream components we first consider a generic stochastic dynamical system that evolves in time *t* according to

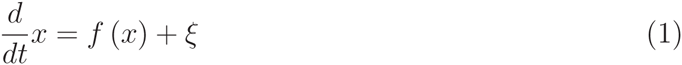

where *x* = (*x*_1_, &, *x_N_*) denotes the variables of the network nodes, *f* describes the intrinsic dynamics of the network, and ξ = (ξ_1;_&, ξ*_Ν_*) is a stochastic external input driving instantaneous state variable fluctuations which carry the information to be routed through the network. We consider a deterministic reference state *x*^(ref)^ (*t*) solving (1) in the absence of signals (ξ = 0).

To quantify how bits of information ’surfing’ on top of such a dynamical state are routed through the network, we use information theoretic measures that quantify the amount of information shared and transferred between nodes, independent of how this information is encoded or decoded. More precisely, we measure information sharing between signal *x_i_* (*t*) and the time *d* lagged signal *x_j_* (*t + d*) of nodes *i* and *j* in the network via the time-delayed mutual information (dMI) [20, 21]

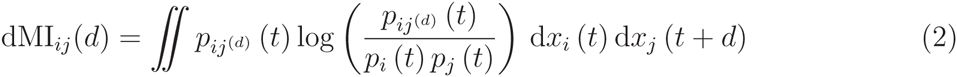

Here *p_i_* (*t*) is the probability distribution of the variable *x_i_* (*t*) of unit *i* at time *t* and *p_i,j(d)_* (*t*) the joint distribution of *x_i_* (*t*) and the variable *x_j_* (*t + d*) lagged by *d*. As a second measure we use the delayed transfer entropy (dTE) [22] (cf. Methods) that genuinely measures information transfer between pairs of units [36]. Asymmetries in the dMI and dTE curves dMI*_ij_* (*d*) and dTE*_i→j_* (*d*) then indicate the dominant direction in which information is shared or transferred between nodes.

To identify the role of the underlying reference dynamical state *x*^(ref)^ (*t*) for network communication a small noise expansion in the signals ξ turns a out to be ideally suited: while the small noise expansion limits the analysis to the vicinity of a specific reference state which is usually regarded as a weakness, in the context of our study, this property is highly advantageous as it directly conditions the calculations on a particular reference state and enables us to extract it’s role for the emergent pattern of information routing within the network. For white noise sources ξ this method yields general expressions for the conditional probabilities *p* (*x* (*t + d*) |*x* (*t*)) that depend on *x*^(ref)^ (*t*). Using this result the expressions for the delayed mutual information (2) and transfer entropy (7) dMI*_i,j_* (*d*) and dTE*_i→j_* (*d*) become a function of the underlying collective reference dynamical state (cf. Methods and Supplementary Section 1). The dependency on this reference state then provides a generic mechanism to change communication in networks by manipulation the underlying collective dynamics. In the following we show how this general principle gives rise to a variety of mechanisms to flexibly change information routing in networks. We focus on oscillatory phenomena widely observed in networks with a communication function [32, 34, 35, 37, 38].

## II. INFORMATION EXCHANGE IN PHASE SIGNALS

Oscillatory synchronization and phase locking [8, 10] provide a natural way for the temporal coordination between communicating units. Key variables in oscillator systems are the phases *ϕ_i_* (*t*) at time *t* of the individual units *i*. In fact, a wide range of oscillating systems display similar phase dynamics [8, 11] (cf. Supplementary Section 2) and phase-based encoding schemes are common, e.g. in the brain [32, 34, 35], genetic circuits [37] and artificial systems [38].

We first focus on systems in a stationary state with a stationary distribution for which the expressions for the dMI and dTE become independent of the starting time *t* and only depend on the lag *d* and reference state *ϕ*^(ref)^ (*t*). To assess the dominant direction of the shared information between two nodes we quantify asymmetries in the dMI curve by using the difference *δ*MI*_i,j_* = MI*_i→j_*·− MI*_i→j_* between the integrated mutual informations 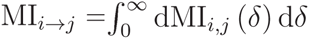 and MI*_i→j_*. If this is positive, information is shared predominantly from unit *i* to *j* while negative values indicate the opposite direction. Analogously, we compute the differences in dTE as *δ*TE*_i,j_*·(cf. Methods and Supplementary Section 3). The set of pairs {*δ*MI*_i,j_*} or {*δ*TE*_i,j_*} for all *i, j* then capture strength and directionality of information routing in the network akin to a functional connectivity analysis in neuroscience [39]. We refer to them as information routing patterns (IRPs).

A range of networks of oscillatory units, with disparate physical interactions, connection topologies and external input signals support multiple IRPs. For instance, in a model of a gene-regulatory network with two oscillatory sub-networks (Fig. 1a) dMI analysis reveals IRPs with different dominant directions (Fig. 1b-d, upper vs. lower sub-panels). The change is triggered by adding an external factor that degrades the transcribed mRNA in one of the oscillators and thereby changes its intrinsic frequency (see Methods). More complex changes in IRPs emerge in larger networks, possibly with modular architecture. In a network of interacting neuronal populations (Fig. 1e) different initial conditions lead to different underlying collective dynamical states. Switching between them induces complicated but specific changes in the IRPs (Fig. 1f-h). Different IRPs also emerge by changing a small number of connections in larger networks. Fig. 1i-l illustrates this for a generic system of coupled oscillators each close to a Hopf bifurcation.

**Figure 1:**
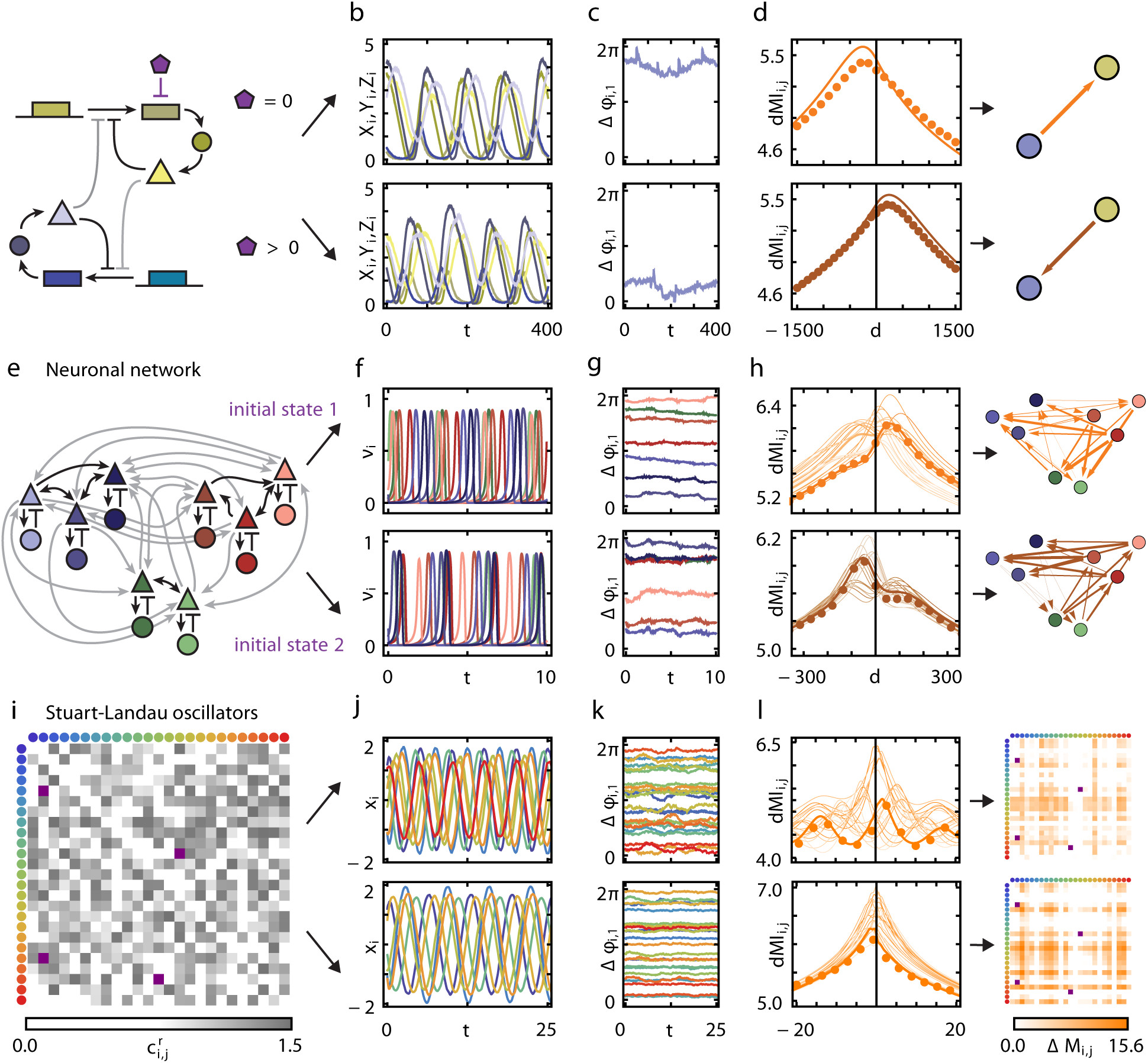
Flexible information routing across oscillatory networks. **a**, Simple model of a gene regulatory network of two coupled biochemical oscillators of Goodwin type (yellow and blue). An additional molecule (purple) degrades the transcribed mRNA in one of the oscillators and thereby changes its intrinsic frequency. Coupling strengths are gray coded (darker color indicates stronger coupling), sharp arrows indicate activating and blunt arrows inhibiting influences. **b**, Stochastic oscillatory dynamics of the system's variables. **c**, Fluctuations of the phases extracted from the full dynamics relative to a reference unit. **d**, Delayed mutual information (dMI_1;2_) between the phase signals. The numerical data (dots) agrees well with the theoretical prediction (4) (solid lines). The asymmetry in the dMI curves around *d* = 0 indicates a directed information sharing pattern summarized in the graphs (right). Arrow thickness indicates the strength of directed information sharing ΔΜI*_i,j_*, measured by the positively rectified differences of the areas below the integrated dMI*_i,j_* (*d*) curve for *d* < 0 and *d* > 0. e–h, Same as in a–d but for a modular network of coupled neuronal sub-populations consisting each of excitatory (triangle) and inhibitory (disk) populations (Wilson-Cowan type dynamics). For the same network two different collective dynamical states accessed by different initial conditions give rise to two different information sharing patterns (f–h top vs. bottom). **i–l**, As in a-d but for generic oscillators close to a Hopf bifurcation (Stuart-Landau oscillators) connected to a larger network. In i and l connectivity matrices are shown instead of graphs. Two different network-wide information routing patterns arise (top vs. bottom in j–l) by changing a small number of connection weights (purple entries in i and l).

In general, several qualitatively different options for modifying network-wide IRPs exist, all of which are relevant in natural and artificial systems: (i) changing the intrinsic properties of individual units (Fig. 1a-d, cf. also Fig. 3a-c below), (ii) modifying the system connectivity (Fig. 1i-l, Fig. 3d-f) and (iii) selecting distinct dynamical states of structurally the same system (Fig. 1e-h, see also Fig. 4 below).

## III. THEORY OF PHASE INFORMATION ROUTING

To reveal how different IRPs arise and how they depend on the network properties and dynamics, we derive analytic expressions for the dMI and dTE between all pairs of oscillators in a network. We determine the phase of each oscillator *i* in isolation by extending its phase description to the full basin of attraction of the stable limit cycle [8, 40]. For weak coupling, the effective phase evolution becomes

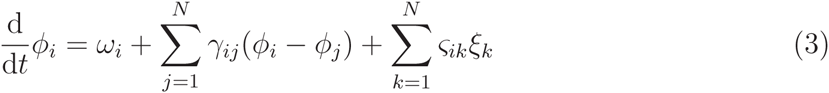

where *ω_i_* is the intrinsic oscillation frequencies of node *i* and the coupling functions *γ_i,j_*(.) depend on the phase differences only. The final sum in (3) models external signals as independent Gaussian white noise processes ξ*_k_* and a covariance matrix ς*_ik_*. The precise forms of *γ_ij_*(.) and ς*_ik_* generally depend on the specific system (Supplementary Section 2).

As visible from Fig. 1e-h, the IRP strongly depends on the underlying collective dynamical state. We therefore decompose the dynamics into a deterministic reference part 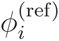 and a fluctuating component 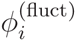. We focus on phase-locked configurations for the deterministic dynamics with constant phase offsets 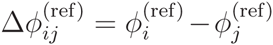. We estimate the stochastic part 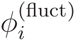 via a small noise expansion (Methods, Supplementary Theorem 1) yielding a first-order approximation for the joint probabilities 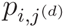. Using (2) together with the periodicity of the phase variables, we obtain the delayed mutual information

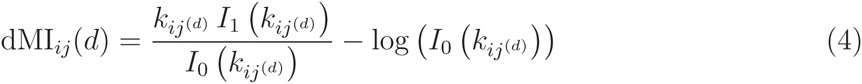

between phase signals in coupled oscillatory networks; here *I_n_* (*k*) is the *n*^th^ modified Bessel function of the first kind, and 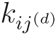 is the inverse variance of a von Mises distributions ansatz for 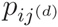. The system’s parameter dependencies, including different inputs, local unit dynamics, coupling functions and interaction topologies are contained in 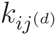. By similar calculations we obtain analytical expressions for dTE*_i→j_* (Methods and Supplementary Theorem 2). Our theoretical predictions well match the numerical estimates (Fig. 1d,h,l, see also Fig. 2c,d below and Supplementary Figs. 1, 2 and 7). For independent input signals (ς*_ik_* = 0 for *i* ≠ *k*) we typically obtain similar IRPs determined either by the delayed mutual information or the transfer entropy (Supplementary Fig. 2).

**Figure 2:**
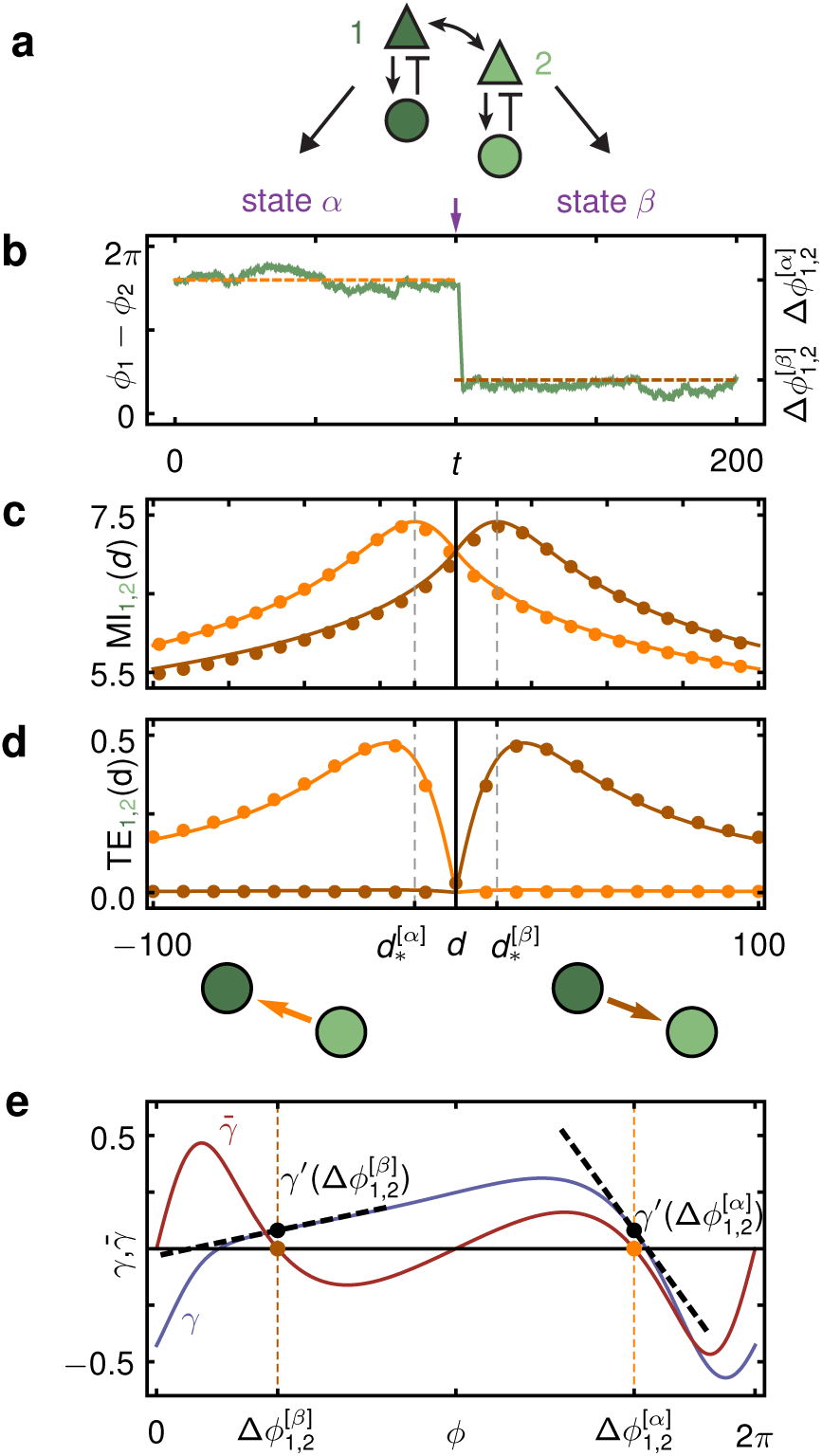
Multi-stable dynamics and flexible anisotropic information routing. **a**, Two identical and symmetrically coupled neuronal circuits of Wilson-Cowan type (dark and light green, modular sub-network in Fg. 1e). **b**, Phase difference Δ*ϕ*_1,2_(*t*):= *ϕ*_1_ (*t*) *− ϕ*_2_ (*t*) between the extracted phases of the two neuronal populations fluctuating around a locked value 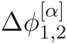 of a stable deterministic collective state *α* (orange); a strong external perturbation (purple arrow) induces a switch to stochastic dynamics around an alternate stable state *β* (brown) with phase difference 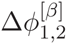. **c**, Delayed mutual information dMI_1,2_ and **d**, transfer entropy dTE_1→2_ between the phase signals in state *α* (orange) and *β* (brown) for numerical data (dots) and theory (lines). The change in peak latencies form 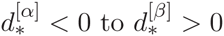 in the dMI_1,2_ and the asymmetry of the dTE_1→2_ curves show anisotropic information routing. Switching between the two dynamical states reverses the information flow pattern (graphs, bottom). **e**, Phase coupling function *γ*(Δ*ϕ*) = γ_12_ (Δ*ϕ*) = γ_21_ (Δ*ϕ*) (blue) and its antisymmetric part 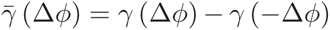 (red). The two zeros of 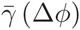 with negative slope indicate the deterministic equilibrium phase differences 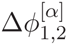 and 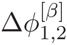 in states *α* and *β*, receptively. The directionality in the information routing pattern arises due to the different slopes of *γ* (Δ*ϕ*) (dashed lines) at the noiseless phase-locking offsets 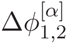 and 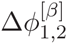.

## IV. MECHANISM OF ANISOTROPIC INFORMATION ROUTING

To better understand how a collective state gives rise to a specific routing pattern with directed information sharing and transfer, consider a network of two symmetrically coupled identical neural population models (Fig. 2a). Due to permutation symmetry, the coupling functions *γ_ij_*, obtained from the phase-reduction of the original Wilson-Cowan-type equations [41] (Methods, Supplementary Section 5), are identical. For biologically plausible parameters this network in the noiseless-limit has two stable phase-locked reference states (*α* and *β*). The fixed phase differences 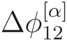 and 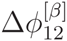 are determined by the zeros of the anti-symmetric coupling 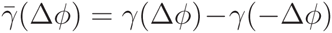 with negative slope (Fig. 2e). For a given level of (sufficiently weak) noise, the system shows fluctuations around either one of these states (Fig. 2b) each giving rise to a different IRP. Sufficiently strong external signals can trigger state switching and thereby effectively invert the dominant communication direction visible from the dMI (Fig. 2c) and even more pronounced from the dTE (Fig. 2d) without changing any structural properties of the network.

The anisotropy in information transfer in the fully symmetric network is due to symmetry broken dynamical states. For independent noise inputs, ς*_ik_* = ς*_i_δ_ik_*, that are moreover small, the evolution of 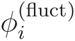, *i* ∈ {1, 2}, near the reference state *α* reduces to

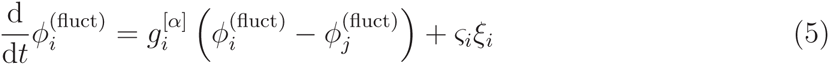

with coupling constants 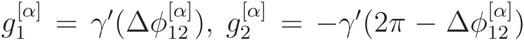 (Methods). As 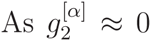 (Fig. 2e), the phase 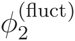 essentially freely fluctuates driven by the noise input ς_2_ξ_2_. This causes the system to deviate from the equilibrium phase difference 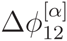. At the same time, the strongly negative coupling 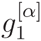 dominates over the noise term ς_1_ξ_1_ and unit 1 is driven to restore the phase-difference by reducing 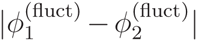. Thus, 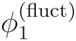 is effectively enslaved to track 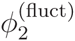 and information is routed from unit 2 to unit 1, reflected in the dMI and dTE curves. The same mechanism accounts for the reversed anisotropy in communication when the system is near state *β* as the roles of unit 1 and 2 are exchanged. Calculating the peak of the dMI curve in this example also provides a time scale 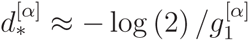 at which maximal information sharing is observed (Methods, Eq. (10)). It furthermore becomes clear that the directionality of the information transfer in general need not be related to the order in which the oscillators phase-lock because the phase-advanced oscillator can either effectively pull the lagging one, or, as in this example, the lagging oscillator can push the leading one to restore the equilibrium phase-difference.

In summary, effective interactions local in state space and controlled by the underlying reference state together with the noise characteristics determine the IRPs of the network. Symmetry broken dynamical states then induce anisotropic and switchable routing patterns without the need to change the physical network structure.

## V. INFORMATION ROUTING IN NETWORKS OF NETWORKS

For networks with modular interaction topology [24–28], our theory relating topology, collective dynamics and IRPs between individual units can be generalized to predict routing between entire modules. Assuming that each sub-network *X* in the noise-less limit has a stable phase-locked reference state, a second phase reduction [23] generalized to stochastic dynamics characterizes each module by a single meta-oscillator with collective phase Φ*_X_* and frequency Ω*_X_*, driven by effective noise sources Ξ*_X_* with covariances Σ*_X_,_Y_*. The collective phase dynamics of a network with *M* modules then satisfies

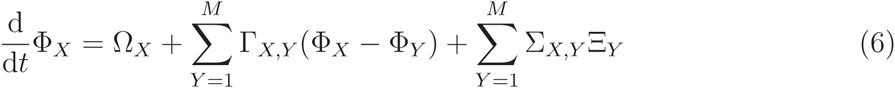

where Γ*_X_*,*_Y_* are the effective inter-community couplings (Supplementary Section 4). The structure of equation (6) is formally identical to equation (3) so that the expressions for inter-node information routing (dMI*_i,j_*, dTE*_i,j_*) can be lifted to expressions on the intercommunity level (dMI*_X_,_Y_*, dTE_X_,_Y_) by replacing node-with community-related quantities (i.e. *ω_i_* with Ω*_X_* or *γ_ik_* with *Γ_XK_*, etc., Supplementary Corollary 3 and 4). Importantly, this process can be further iterated to networks of networks, etc. Fig. 3 shows examples of information flow patterns resolved at two scales. The information routing direction on the larger scale reflects the majority and relative strengths of IRPs on the finer scale.

**Figure 3:**
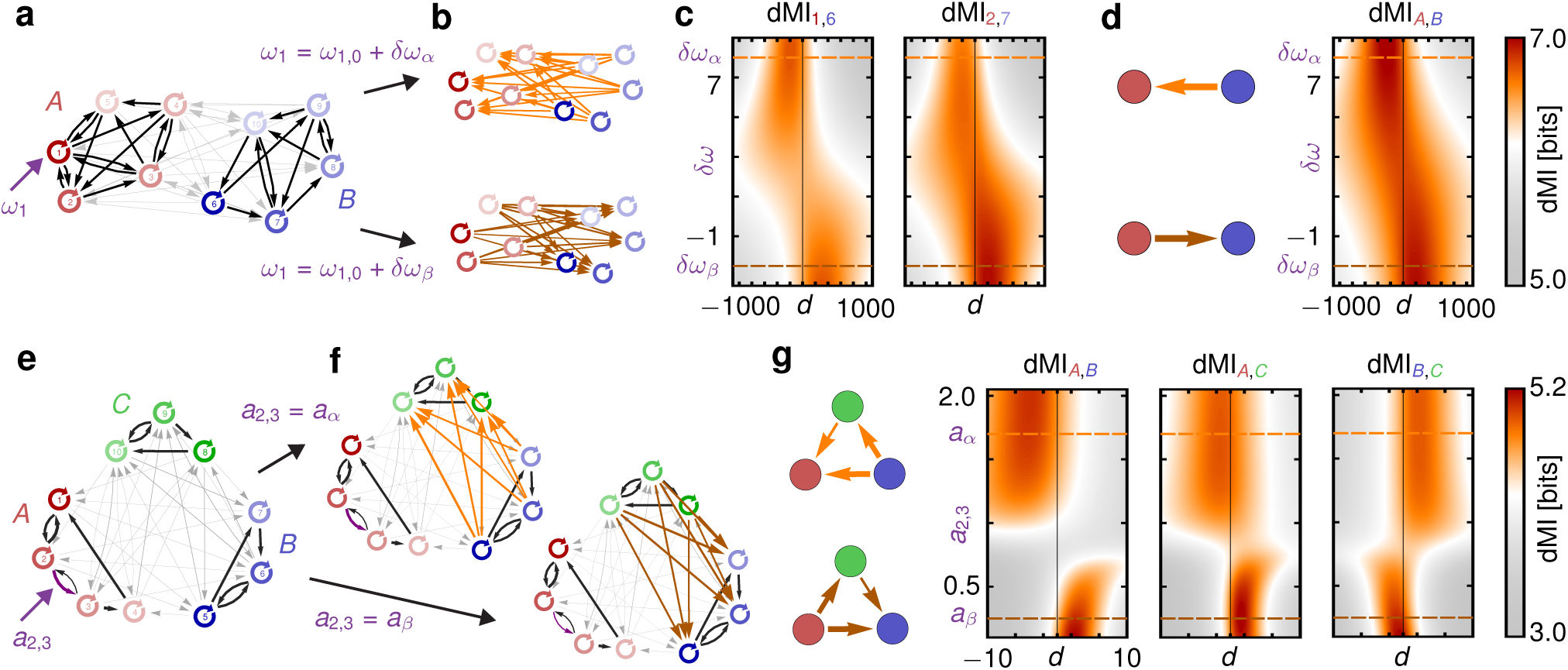
Remote rerouting of information in modular networks: Local changes trigger global information rerouting. **a**, Network with two coupled communities *A* and *B* (red and blue) of oscillators close to a Hopf bifurcation. Changing the intrinsic frequency of a single node *i* = 1 from *ω_1_* + *δω_α_* to *ω_1_* + *δω_β_* induces a collective reorganization of equilibrium phase differences, that result in **b**, oppositely directed information sharing patterns (top vs bottom). **c**, dMI between two pairs of nodes from the two different clusters as a function of the time delay *d* and frequency change *δω*_1_ of oscillator 1. **d**, Information flow patterns calculated from the hierarchically reduced system for the two configurations in b (left) and as a function of *δω*_1_ (right) reflect the inversion of the IRPs on the finer scale (b,c). **e**, Network of three coupled modules of phase oscillators. **f**, A change in the connection strength *a*_2_,_3_ from *a_α_* to *a_β_* between two nodes (3*_A_* → 2*_a_*) in sub-network *A* induces an inversion of information routing direction between the remote sub-networks *B* and *C*. **g**, Full information routing patterns calculated form the hierarchical reduced system for *a*_2_,_3_ = *a_α_* and *a*_2,3_ = *a_α_* (left) and as a function of *a*_2_,_3_ for all pairs of modules (density plots, right). The transition is not continuous but rather switch like.

## VI. NON-LOCAL INFORMATION REROUTING VIA LOCAL INTERVENTIONS

The collective quantities in the system (6) are intricate functions of the network properties at the lower scales. Intriguingly, the coupling functions Γ*_X,Y_* not only depend on the non-local interactions γ*_iX jY_* between units *i_X_* of module *X* and *j_Y_* of cluster *Y* but also on purely local properties of the individual clusters. In particular, the form of Γ*_X,Y_* is a function of the intrinsic local dynamical states 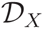 and 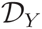 of both clusters as well as the phase response *Z_X_* of sub-network *X* (see Methods and Supplementary Section 4). Thus IRPs on the entire network level depend on local community properties. This establishes several generic mechanisms to globally change information routing in networks via local changes in modular properties, local connectivity, or via switching of local dynamical states.

In a network consisting of two sub-networks (Fig. 3a) the local change of the frequency of a single Hopf-oscillator in sub-network *A* induces a non-local inversion of the information routing between cluster *A* and *B* (Fig. 3b-d). In Fig. 3e-f the direction in which information is routed between two sub-networks *B* and *C* of coupled phase oscillators is remotely changed by increasing the strength of a local link in module *A*. The origin in both examples is a non-trivial combination of several factors: The (small) manipulations alter the collective cluster frequency Ω*_Α_*, the local dynamical state 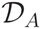 which in turn change the collective phase response *Z_A_* and the effective noise strength Ξ*_Α_* of cluster *A* (Supplementary Fig. 3). These changes all contribute to changes in the effective couplings Γ*_X,Y_* as well as in the inter-cluster phase-locking values ΔΦ*_X,Y_* = Φ*_X_* − Φ*_Y_*. Taken together this causes the observed inversions in information routing direction. Interestingly, the transition in information routing has a switch like dependency on the changed parameter (Fig. 3c,d,g) promoting digital-like changes of communication modes.

## VII. COMBINATORIAL INFORMATION ROUTING PATTERNS

As an alternative to interventions on local properties, also switching between multi-stable local dynamical states 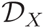 can induce global information rerouting. In the example in Fig. 4, each of the *M* = 3 modules *X ∈* {*A*, *B, C*} exhibits 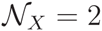 alternative phase-locked states (labeled *α_X_* and *β_X_*, Supplementary Section 5.1). For sufficiently weak coupling, this local multi-stability is preserved in the dynamics of the entire modular network. Consequently each choice of the 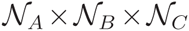 possible combinations of “local” states gives rise to at least one network-wide collective state. Certain combinations of local states can give rise to one or even multiple globally phase-locked states (e.g. [*α_A_β_Β_α_C_*] in Fig. 4). Others support non-phase locked dynamics that gives rise to time-dependent IRPs (cf. Fig. 4c and below). Thus. varying local dynamical states in a hierarchical network flexibly produces a combinatorial number 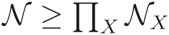 of different IRPs in the same physical network.

**Figure 4:**
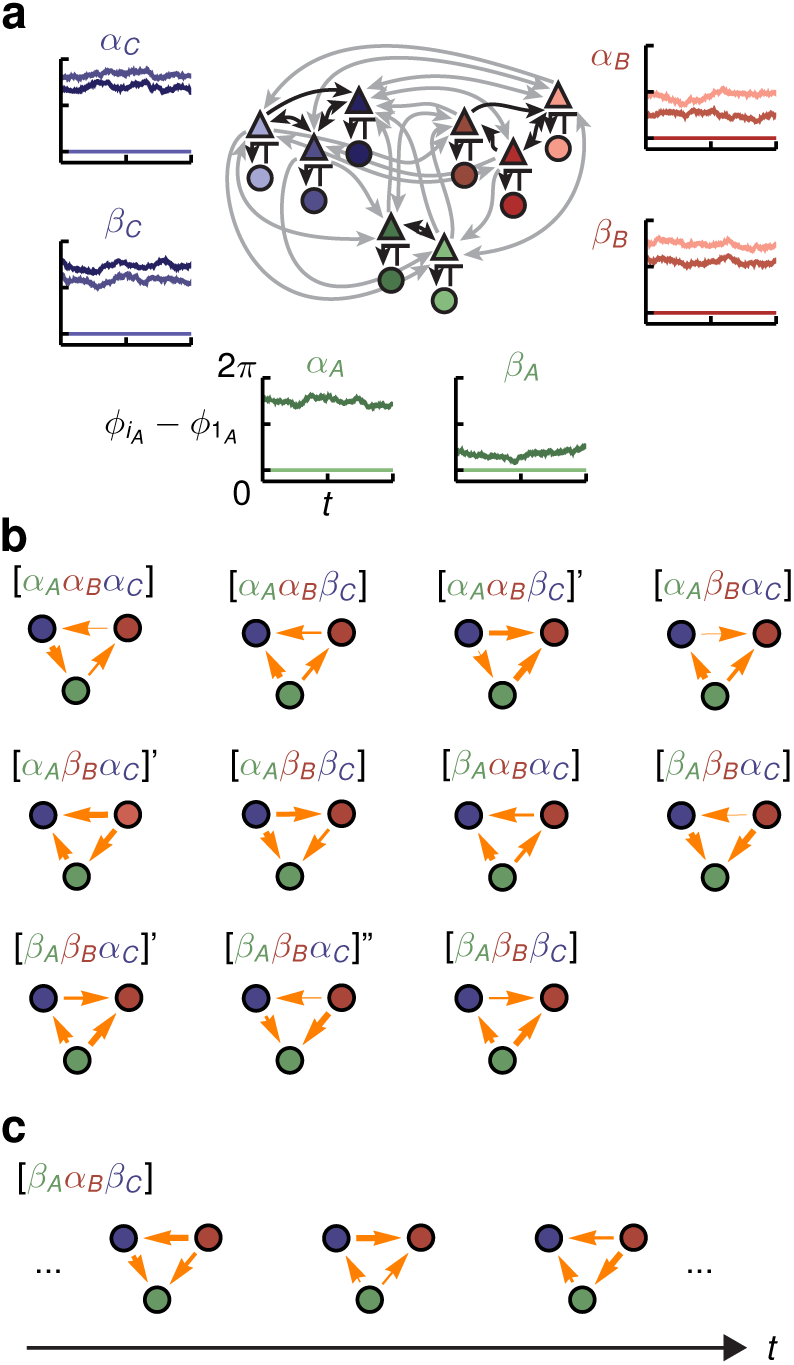
Switching between combinatorially many information routing patterns. **a**, Modular circuit as in Fig. 1e. Without inter-module coupling, each of the *M* = 3 communities *X ∈* {*A, B,C*} exhibits multi-stability between two phase-locked configurations, denoted as states *α_X_* and *β_X_* (insets). **b**, Information routing patterns between the hierarchically reduced sub-networks for different combinations of the local dynamical states 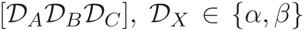 that give rise to globally phase-locked dynamics. Arrows between nodes *X* and *Y* indicate strength (line width) and sign (arrow direction) of the difference in integrated dTE curves between the nodes. The same local dynamical configuration 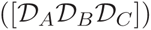 can give rise to more than one globally locked collective state marked with dashes, i.e. 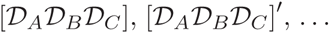. **c**, The local dynamical state configuration [*β_Α_α_Β_β_C_*] generates a periodic global dynamical state (cf. Supplementary Figure 8) in which the hierarchically reduced information routing pattern (graphs) becomes time-dependent (cf. also Supplementary Section 6).

## VIII. TIME-DEPENDENT INFORMATION ROUTING

General reference states, including periodic or transient dynamics, are not stationary and hence the expressions for the dMI and dTE become dependent on time *t*. For example, Fig. 4c shows IRPs that undergo cyclic changes due to an underlying periodic reference state (cf. also Supplementary Figure 8a-c). In systems with a global fixed point systematic displacements to different starting positions in state space give rise to different stochastic transients with different and time-dependent IRPs (Supplementary Figure 8d). Similarly, switching dynamics along heteroclinic orbits constitute another way of generating specific progressions of reference dynamics. Thus information ’surfing’ on top of non-stationary reference dynamical configurations naturally yield temporally structured sequences of IRPs, resolvable also by other measures of instantaneous information flow, e.g. [36, 42, 43].

## IX. DISCUSSION

The above results establish a theoretical basis for the emergence of information routing capabilities in complex networks when signals are communicated on top of collective reference states. We show how information sharing (dMI) and transfer (dTE) emerge through the joint action of local unit features, global interaction topology and choice of the collective dynamical state. We find that information routing patterns self-organize according to general principles (cf. Figs. 2, 3, 4) and can thus be systematically manipulated. Employing formal identity of our approach at every scale in oscillatory modular networks (Eq. (3) vs. (6)) we identify local paradigms that are capable of regulating information routing at the non-local level across the whole network (Figs. 3, 4).

In contrast to self-organized technological routing protocols where local nodes use local routing information to locally propagate signals, such as in peer-to-peer networks [44], in the mechanism studied here the information routing modality is set by the entire network’s collective dynamics. This collective reference state typically evolves on a slower time scale than the information carrying fluctuations that surf on top of it and is thus different from signal propagation in cascades [45] or avalanches [46] that dominate on shorter time scales.

We derived theoretical results based on information sharing and transfer obtained via delayed mutual information and transfer entropy curves. Using these abstract measures our results are independent of any particular implementation of a communication protocol and thus generically demonstrate how collective dynamics can have a functional role in information routing. For example, in the network in Fig. 2 externally injected streams of information are automatically encoded in fluctuations of the rotation frequency of the individual oscillators. The injected signals are then transmitted through the network and decodable from the fluctuating phase velocity of a target unit precisely along those pathways predicted by the current state-dependent IRP (Supplementary Section 7).

Our theory is based on a small noise approximation that conditions the analysis onto a specific underlying dynamical state. In this way we extracted the precise role of such a reference state for the network’s information routing abilities. For larger signal amplitudes or in highly recurrent networks in which higher-order interactions can play an important role the expansion can be carried out systematically to higher orders using diagrammatic approaches [47] or numerically to accounting for better accuracy and non-Gaussian correlations (cf. also Supplementary Section 3.4).

In systems with multi-stable states two signal types need to be discriminated: those that encode the information to be routed and those that indicate a switch in the reference dynamics and consequently the IRPs. If the second type of stimuli is amplified appropriately a switch between multi-stable states can be induced that moves the network into the appropriate IRP state for the signals that follow. For example, in the network of Fig. 2 a switch from state *α* to *β* can be induced by a strong positive pulse to oscillator 2 (and vice versa). If such pulses are part of the input a switch to the appropriate IRP state will automatically be triggered and the network auto-regulates its IRP function. More generally a separate part of the network that effectively filters out relevant signatures indicating the need for a different IRP could provide such pulses. Moreover, using the fact that local interventions are capable to switch IRPs in the network the outcomes of local computations can be used to trigger changes in the global information routing and thereby enable context-dependent processing in a self-organized way.

When information surfs on top of dynamical reference states the control of IRPs is shifted towards controlling collective network dynamics making methods from control theory of dynamical systems available to the control of information routing. For example, changing the interaction function in coupled oscillators systems [18] or providing control signals to a subset of nodes [48, 57] are capable of manipulating the network dynamics. Moreover, switch like changes (cf. Fig. 3) can be triggered by crossing bifurcation points and the control of information routing patterns then gets linked to bifurcation theory of network dynamical systems.

While the mathematical part of our analysis focused on phase signals, including additional amplitude degrees of freedom into the theoretical framework can help to explore neural or cell signaling codes that simultaneously use activity- and phase-based representations to convey information [49]. Moreover, separating IRP generation, e.g. via phase configurations, from actual information transfer, for instance in amplitude degrees of freedom, might be useful for the design of systems with a flexible communication function.

The predicted phenomena, including non-local changes of information routing by local interventions, could be directly experimentally verified using methods available to date, such as electrochemical arrays [18] or synthetic gene regulatory networks [5] (Supplementary Section 5.3). In addition our results are applicable to the inverse problem: Unknown network characteristics may be inferred by fitting theoretical expected dMI and dTE patterns to experimentally observed data. For example, inferring state-dependent coupling strengths could further the analysis of neuronal dynamics during context-dependent processing [33, 35, 39, 50–52].

Modifying inputs, initial conditions or system-intrinsic properties may well be viable in many biological and artificial systems whose function requires particular information routing. For instance, on long time scales, evolutionary pressure may select a particular information routing pattern by biasing a particular collective state in gene regulatory and cell signaling networks [2, 15, 53]; on intermediate time scales, local changes in neuronal responses due to adaptation or varying synaptic coupling strength during learning processes [13] can impact information routing paths in entire neuronal circuits; on fast time scales, defined control inputs to biological networks or engineered communication systems that switch the underlying collective state, can dynamically modulate information routing patterns without any physical change to the network.

**Methods**: *Transfer Entropy.* The delayed transfer entropy (dTE) [22] from a time-series *x_i_*(*t*) to a time-series *x_j_*(*t*) is defined as

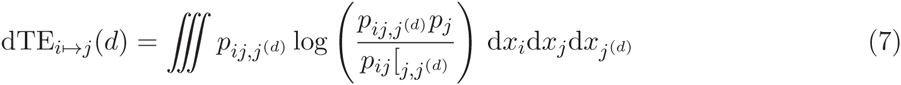

with joint probability 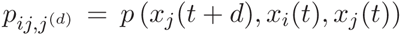. This expression is not invariant under permutation of *i* and *j*, implying the directionality of TE. For a more direct comparison with dMI in Figure 2, we define dTE_ij_(*d*) by dTE_i→j_·(*d*) for *d* > 0 and by dTE*_j→i_*(−*d*) for *d* < 0.

*Dynamic Information routing via dynamical states.* For a dynamical system (1) the reference deterministic solution *x*^(ref)^ (*t + s*) starting at *x* (*t*) is given by the deterministic flow 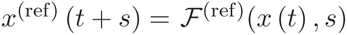. The small noise approximation for white noise ξ then yields

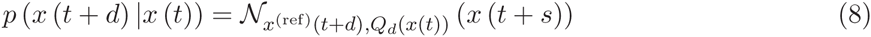

where 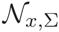 denotes the normal distribution with mean *x* and covariance matrix Σ, 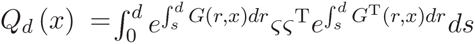 and 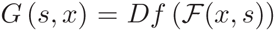. From this and the initial distribution *p* (*x*(*t*)) the delayed mutual information and transfer entropy dMI*_i,j_* (*d, t*) and dTE*_i→j_* (*d, t*) are obtained via (2) and (7). The result depends on time *t*, lag*d* and the reference state *x*^(ref)^.

*Oscillator Networks.* In Fig. 1a, we consider a network of two coupled biochemical Goodwin oscillators [14, 54]. Oscillations in the expression levels of the molecular products arise due to a nonlinear repressive feedback loop in successive transcription, translation and catalytic reactions. The oscillators are coupled via mutual repression of the translation process [55]. In addition, in one oscillator changes in concentration of an external enzyme regulate the speed of degradation of mRNAs, thus affecting the translation reaction, and, ultimately, the oscillation frequency. In Fig. 1e, 2, 4 we consider networks of Wilson-Cowan type neural masses (population signals) [41]. Each neural mass intrinsically oscillates due to antagonistic interactions between local excitatory and inhibitory populations. Different neural masses interact, within and between communities, via excitatory synapses. In the generic networks in Fig. 1i and Fig. 3a each unit is modeled by the normal form of a Hopf-bifurcation in the oscillatory regime together with linear coupling. Finally, the modular networks analyzed in Figures 3a and 3b are directly cast as phase-reduced models with freely chosen coupling functions. See the Supplementary Information for additional details, model equations and parameters and phase estimation.

*Analytic derivation of the* dMI *and* dTE *curves.* In the small noise expansion [56], both dMI and dTE curves have an analytic approximation: For stochastic fluctuations around some phase-locked collective state with constant reference phase offsets Δ*ϕ_ij_*, = *ϕ_i_* − *ϕ_j_* the phases evolve as 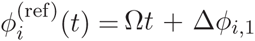 in the deterministic limit, where Ω = *ω_i_ +* Σ*_k_ γ_ik_* (Δ*ϕ_ik_*) is the collective network frequency and the *γ_ij_* (.) are the coupling functions from Eq. (3). In presence of noise, the phase dynamics have stochastic components 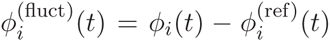. In first order approximation, independent noise inputs *ς_ij_* = ς*_i_δ_ij_* yield coupled Ornstein-Uhlenbeck processes

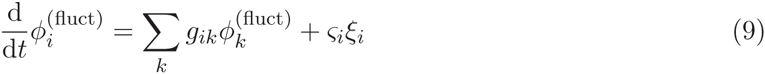

with linearized, state-dependent couplings given by the Laplacian matrix entries 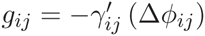 and 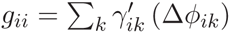. The analytic solution to the stochastic equations (9) provides an estimate of the probability distributions, *P_i_*, 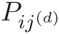 and 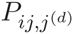. Via (2) this results in a prediction for dMI*_ij_* (*d*), Eq. (4), as a function of the matrix elements 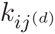 specifying the inverse variance of a von Mises distribution ansatz for 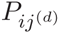. Similarly via (7) an expression for dTE*_i→j_*(*d*) is obtained. For the dependency of 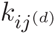 and dTE*_i→j_*(d) on network parameters and further details, see the derivation of the Theorems 1 and 2 in the Supplementary Information.

*Time scale for information sharing.* For a network of two oscillators as in Fig. (2) with linearized coupling strengths 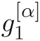 and 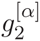 and 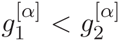, maximizing dMI_12_ (*d*) (see Supplementary Information for full analytic expressions of dMI and dTE in two oscillator networks) yields

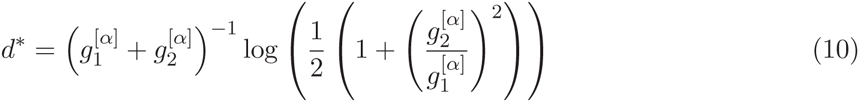

*Collective phase reduction.* Suppose that each node *i* = *i_X_* belongs to a specific network module *X* out of *M* ≤ *N* non-overlapping modules of a network. Then equation (3) can be simplified to (6) under the assumption that in the absence of noise every community *X* has a stable internally phase-locked state 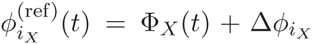, where 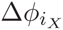 are constant phase offsets of individual nodes *i_X_*. Every community can then be regarded as a single meta-oscillator with a collective phase Φ*_X_*(*t*) and a collective frequency 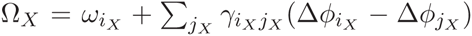. The vector components of the collective phase response *Z_X_*, the effective couplings Γ*_XY_* and the noise parameters Σ*_XY_* and Ξ*_X_* are obtained through collective phase reduction and depend on the respective quantities 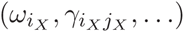 on the single-unit scale (see Supplementary Section 4 for a full derivation).

## Acknowledgements

We thank T. Geisel for valuable discussions. Partially supported by the Federal Ministry for Education and Research (BMBF) under grants no. 01GQ1005B [CK, DB, MT] and 03SF0472E [MT], by the NVIDIA Corp., Santa Clara, USA [MT], a grant by the Max Planck Society [MT], by the FP7 Marie Curie career development fellowship IEF 330792 (DynViB) [DB] and an independent postdoctoral fellowship by the Rockefeller University, New York, USA [CK].

## Author contributions

All authors designed research. C.K. derived the theoretical results, developed analysis tools and carried out the numerical experiments. All authors analyzed and interpreted the results and wrote the manuscript.

## Additional information

The authors declare no competing financial interests. Supplementary information accompanies this paper. Correspondence and requests should be addressed to CK.

